# Serum free expansion and transduction of human V gamma 9-V delta 2 T cells for adoptive immunotherapy

**DOI:** 10.64898/2026.01.08.698334

**Authors:** Thamizhselvi Ganapathy, Muthuganesh Muthuvel, Augustine Thambaiah Prabakumar, Seth Sakshi, Aleya Tabasum, Lalitha Priyadharshini Sivamani, Mohankumar Murugesan, Sunil S Raikar, Trent Spencer, Aby Abraham, Alok Srivastava, Sunil Martin

**Affiliations:** Centre for Stem Cell Research (a unit of inStem, Bengaluru), Christian Medical College Campus, Vellore 632002, Tamil Nadu, India; Department of Hematology, Christian Medical College Vellore, Ranipet Campus – 632517, Tamil Nadu, India; Department of Pediatrics and Aflac Cancer and Blood Disorders Center, Emory University School of Medicine, Atlanta, GA, USA; Synthetic Immunology Laboratory, Cancer Research Division, Rajiv Gandhi Center for Biotechnology, Thiruvananthapuram 695 011, Kerala, India; Manipal Academy of Higher Education, Manipal, 576104, Karnataka, India

**Keywords:** V gamma 9-V delta 2 T cells, Zoledronic Acid, Cancer Immunotherapy, Off-the-shelf, Serum Free Media

## Abstract

**Background and Aim:** Gamma delta T cells are multivalent immune cells with innate and adaptive features that sense a broad spectrum of tumor-associated stress patterns to lyse them without prior potentiation. They are therefore a good option for developing an off-the-shelf therapeutic immune cell product with antitumor functions. Therefore, we aimed to develop an optimized cGMP compatible protocol for expanding V gamma 9-V delta 2 T cells in a serum-free media for adoptive immunotherapy applications.

**Methods and Results:** Peripheral blood mononuclear cells (PBMC) from healthy donors were activated with Zoledronic Acid (ZOL) and IL-2 for 14 days. Cell proliferation, fold expansion, and phenotype were monitored. Among the regular donors, the baseline levels (day 0) of CD3+Vgamma9+delta2+ T cells were (05.10+/−0.74%). There was a robust expansion of V gamma 9-V delta 2 T cells in the serum-free media (110+/−29.89-fold). During processing, an abrupt reduction of αβ T cells was observed as early as day +07. After two weeks, 87.82+/−5.11% (*n=8*) of T cells were CD3+Vgamma9+ in Optimizer with 97.15+/−0.7% of the CD3+Vgamma9+ T cells positive for delta2. The CD3+Vgamma9+ NKG2D+ increased during expansion, reaching 93.76+/−1.55% expression on day +14. V gamma 9-V delta 2 T cells expanded in the serum-free (Optimizer) media had relatively reduced variance and reduced over all yield and fold expansion but comparable percentage of Vgamma9+ and αβ TCR+ T cells on day 14. A flow cytometry-based tumor-toxicity assay gauged the antitumor functions against K562 cell lines. Pretreatment of K562 cells with ZOL differentially enhanced (2.48+/−0.76 fold) the cytotoxic capacity of V gamma 9-V delta 2 T cells in a donor-dependent manner. The conditions for lentiviral transduction and transgene expressions of V gamma 9-V delta 2 T cells were improvised as gauged by GFP expression driven by CMV promoter (39.09+/− 8.94%) without compromising the viability.

**Conclusion:** We have optimized a cGMP compatible protocol for expansion of human V gamma 9-V delta 2 T cells in a serum-free media with high purity and viability in as early as 07 days. Pretreatment of target tumor cells with ZOL enhanced the cytotoxicity, revealing the impact of V gamma 9-V delta 2 T cell intrinsic factors in tumor lysis. This protocol, may be scaled up for clinical translation for adoptive immunotherapy.

## Introduction

Promising clinical results in otherwise refractory malignancies triggered extensive clinical investigations in adoptive immunotherapy[1]. Most clinical trials focused on CD8+ and CD4+ T cells with αβ TCRs, which specifically recognize the tumor antigens presented as peptide-HLA to mount a durable antitumor immune response[2]. The success of αβ T cells in adoptive immunotherapy is owing to 1) well-established expansion and genetic engineering protocol, 2) viability and functionality in the tumor microenvironment, and 3) the ability to confer long-term immunity due to antigen-specific memory response[3]. However, in specific patient subsets, the adoptive immunotherapy of multiple hematological and solid tumors poses fresh challenges due to quality of autologous T cells, antigen escape and cytokine toxicity often associated with HLA-restricted αβ T cells. Moreover, graft versus host disease (GvHD) due to HLA-TCR mismatch of allogenic T cells limits their applicability in generating ‘off-the-shelf’ products[4]. Remarkably, the untenable price tag of the cell therapeutics limit patient access in the current custom generated, autologous models of immune cell therapy. Batch production of ‘off-the-shelf’ immune cells is like to make batch produced product more affordable and may enable broader adoption further reducing the cost. The complicated logistics of immune cell therapy and time lag between blood collection and CAR T cell infusion can worsen disease condition and alter the patient may fall out of eligibility criteria at the time of infusion[5].

gamma delta T cells are unique subsets of T cells combining innate and adaptive features that sense ‘tumor patterns’ than tumor-specific antigens in an HLA-independent manner[6]. They represent a small fraction (1-5%) of the T cell compartment and display a specific distribution. Unlike their αβ counterpart, gamma and delta chains selected by V(D)J recombination are utilized in the TCR heterodimeric complex, *albeit* with minimal repertoire[6]. V gamma 9-V delta 2 T cells - a subset abundant in the peripheral blood - recognize phospho-antigens elevated in the malignant or infected cells[7]. Like NK cells, gamma delta T cells also detect tumors by a constellation of ‘inhibitory’ and ‘activating’ receptors, thus detecting MHC downregulated and sub-optimally immunogenic tumors with reduced mutation load. Although lymphoid in origin, gamma delta T cells retain several features of the innate immune cells, such as expression of high-affinity Fc binding region and pattern recognition receptors apart from its ability to participate in antigen presentation[8]. This ‘multivalent immunity’ of gamma delta T cells makes it a desirable candidate for adoptive immunotherapy applications. gamma delta T cells therapy had a very high safety profile with no indication for progressive disease. A recent study reported complete remission of B-cell Non-Hodgkin Lymphoma patients with **‘**richter transformation**’** following Haploidentical gamma delta T cel therapy[9]. gamma delta T cells are therefore a promising modality in such unique and clinically challenging situations.

A remarkable limitation in the clinical scale expansion of V gamma 9-V delta 2 T cells is the xenogeneic serum used in the media, which necessitates additional downstream processing apart from risks of anti-xenogenic immune reaction and adventitious microbes further escalating the risk[10]. With increasing requirements for components of immune cell therapy, serum supply may not meet the demand, let alone the animal ethics aspects of the serum collection. Moreover, serum also contributes to the variabilities and inconsistencies in the expansion and function of the cell therapeutics[11].

Depletion of αβ T cells is a customary step in the allogenic gamma delta T cell products due to concerns of outgrowing allogenic αβ T cells with potential for graft versus host disease (GvHD)[12]. Depletion of αβ T cells by anti-TCR αβ antibody, six days after gamma delta T cell activation in the culture is reported to remarkably enhance the gamma delta T cells by day 14[13]. IL15 is a homeostatic cytokine required for the proliferation of NK and gamma delta T cells with an inhibitory reaction on effector αβ T cells[14].

Globally, immune cell therapy is making its way as a standard of care in blood malignancies. Developing a robust protocol that affordably generates a clinically relevant number of therapeutic immune cells is one of the primary goals for immunotherapy [15]. Given the versatile immunity of gamma delta T cells to target infected and malignant cells, we optimized a protocol to expand viable and functional V gamma 9-V delta 2 T cells in and serum-free conditions. This protocol can be modified for manual or automated production of indigenous V gamma 9-V delta 2 T cells in open or closed systems for adoptive immunotherapy.

## Methodology

### Cells and reagents

Peripheral blood mononuclear cells (PBMCs) were isolated from the peripheral blood of healthy volunteers after obtaining informed consent. K562 cells (*Courtesy* Spencer Lab, Emory University and R.V. Shaji, CSCR/CMC, Vellore) were transduced with pTS-GFP lentiviral vector and sorted to generate a K562-GFP cell line 15 days after the transduction. The following reagents were used for various experiments: Zoledronic acid (Zoledra, Axiommax Oncology), recombinant human IL-2 (rIL-2, Miltenyi Biotec), 7-AAD as well as antibodies against Vgamma9 and CD27 (BD Biosciences). TCRαβ, Vdelta2, CD3 and CD45 (Biolegend).

### Expansion of V gamma 9-V delta 2 T cells

V gamma 9-V delta 2 T cells were expanded from PBMCs in the presence of ZOL 5μM) and rhIL-2 (1000IU) on RPMI 1640 with 10% Fetal bovine serum with 1% PenStrep, 1% HEPES and 1% GluMAX. Media was changed 03 days after stimulation and every 2 days until day 14. Except for the experiments comparing serum and cytokines, the cells were washed while changing the media. Cell proliferation and viable cell count were monitored by Countess (Thermofisher Scientific). The diagrammatic workflow for the project is illustrated here (**Suppl. Fig.01**).

#### Lentiviral vector production

Pseudoviral particle generation was performed as reported previously[16]. The methods to develop a 3^rd^ generation Self-Inactivating (SIN) vector with a superior safety profile were described previously[17]. After sequence authentication, the transfer plasmid was transfected into HEK 293T cells along with the packaging vector and envelope protein of vesicular stomatitis virus (VSV-G)-expressing vector in the presence of 2.5M CaCl2. The supernatant was collected 48 hrs after the transduction, and lentivirus was concentrated (Lenti-X concentrator kit, Takara) and was titered by flow cytometry as previously cited[18]. 0.5e6 of V gamma 9-V delta 2 T cells (day 07) were plated and transduced in the presence of pseudoviral particles at an MOI of 80. The cells were washed 2 days after the transduction and maintained for another two days before flow cytometry.

#### Phenotyping by flow cytometry and data analysis

V gamma 9-V delta 2 T cell cultures were monitored on day 0, day 7, and day 14. A viable cell was selected by excluding 7-AAD positive cells, which in turn was used for calculating percentage viability. The αβ and gamma delta T Cells were stained on day 0, day 7, and day 14 of culture, whereas the non-lineage markers were monitored on day 0 and day 14. The cells were stained with anti-CD3, anti-TCR αβ, anti-TCR gamma delta, anti-CD19 (B cell marker), anti CD56-(NK cell marker), anti CD14-(monocyte marker), anti CD69 and NKG2D (Biolegend). All antibodies are raised against the human antigens. These antibodies are added sequentially in an aliquot of cultured cells washed and resuspended in FACS buffer (PBS+0.5% FBS) and blocked in Fc block (Miltenyi) for 15 minutes at 4°C. The antibodies were incubated on ice for 20 minutes. The cells were washed in FACS buffer 2X at 300Xg for 10 minutes. The cells were then stained with 7-AAD (BD Pharmingen) to segregate live cells from dead cells. The cells were acquired by BD FACS Celesta and analysed by *FlowJo v10.9* software. The cells were gated on the lymphocyte-monocyte gate, and subsequently from the 7-AAD (−) CD3+ live gate αβ and gamma delta T cell populations were analysed. The non-lineage markers were analysed from the 7-AAD^neg^CD3^neg^ gate (**Fig.01**).

**Fig. 01:**
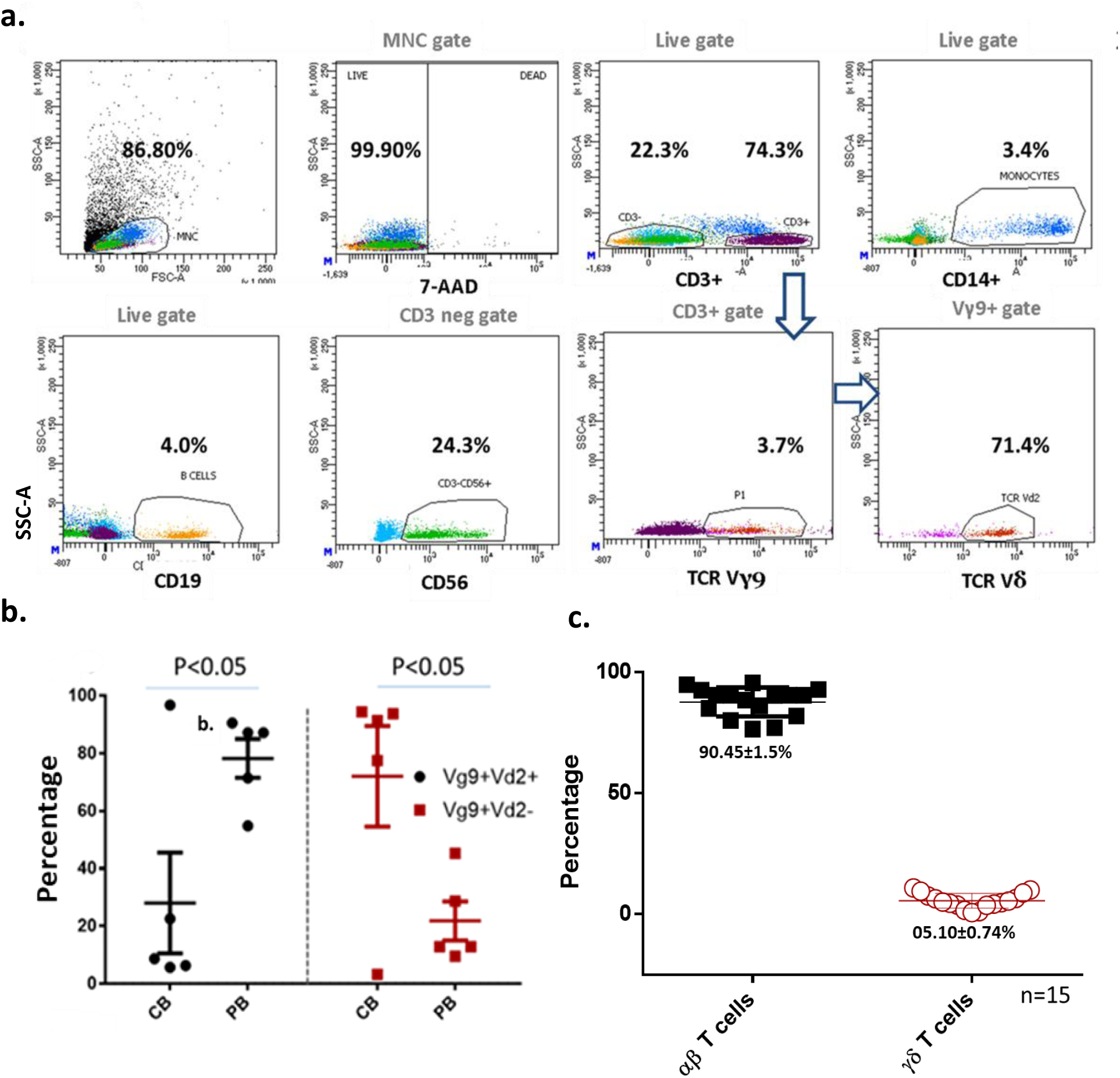
Baseline levels of gamma delta and αβ T cells in regular donors: **a)** The gating strategy for assessing the T cell and non-lineage subsets. CD14+ myelo-monocytic population and CD19+ B lineage populations were assessed within this live gate. The αβ and gamma9 positive population were analysed within the CD3+ gate and delta2 + populations within the gamma9 gate. **b)** The percentage expression of gamma9(+) delta2(+) and gamma9(−) delta(−) cells, n=5 **c)** The baseline expression levels of CD3(+)gamma9(+) T cells as percentage of PBMCs (mean+/−sem) across the normal donor samples, n=15. The statistical significate is estimated by Student’s T test. * P<0.05, **p<0.00, ***p<0.001.n.s is not significant.

### Cytotoxicity assay

A flow cytometry-based cytotoxicity assay was optimized to gauge the tumor toxicity of expanded gamma delta T cells. Briefly, the K562-GFP effector cells were co-cultured with gamma delta T cells at 1:1,1:5 ratio for 4Hrs with effector alone and target alone as controls. GFP positive and 7-AAD positive cells were monitored by flow cytometry at the beginning of the experiment (0^th^ Hr) and 4^th^ Hr. The percentage of death was calculated from GFP+ and 7-AAD+ population. The percentage cytotoxicity was computed by deducing the percentage of cell death at 0^th^ Hr from the percentage of cell death at 4^th^ Hr.

### Statistics

The protocol for the project was approved by the institutional review board and institutional biosafety committee of Christian Medical College Vellore. Each experiment is repeated with the indicated number of normal donors and replicates. The data sets were statistically analyzed using the student’s t-test. Statistical analyses were performed with GraphPad Prism 7. Unless otherwise stated, data is represented as mean +/− standard error mean. Two-tailed unpaired t-tests were used to analyze normally distributed data with two groups. Significance is indicated by *p<0.5, **P<0.01, ***p<0.001, ****p<0.0001.

## Results

### Peripheral blood of healthy volunteers as a source for V gamma 9-V delta 2 T cells

Mononuclear cells derived from cord blood and peripheral blood are the primary sources of gamma delta T cells. Since delta2(**+**) T cells have natural antitumor functions without prior potentiation, we screened PBMCs from healthy volunteers to quantify the percentage of delta2+ T cells in the Vgamma9 compartment. The 7-AAD negative and live cells within the lymphocyte-monocyte gate were examined for CD14+monocytes, CD19 (+) B lineage cells, CD3 neg CD56 (+) NK cells, CD3 (+) Vgamma9 TCR (+) T cells and CD3 (+) αβ TCR (+) T cells. The fractions other than CD3 (+) Vgamma9 TCR (+) T cells were designated as non-lineage cells. More than 70% of the Vgamma9(+) T cells are positive for delta2 at baseline (**Fig.1a**). The V gamma 9-V delta 2 T cells were found to be higher in the peripheral blood whereas delta2^neg^ cells dominated the cord blood mononuclear cells (72.05+/−17.45%) than Vgamma9+delta2+ cells (27.98+/−17.45%) (**Fig.1b**). Hence, the V gamma 9-V delta 2 T cells from the peripheral blood were chosen for optimizing the expansion conditions. The baseline levels of CD3+Vgamma9+ T cells were 05.10+/−0.74% across 15 healthy donors, whereas 90.45+/−1.5% of CD3+ were αβ T cells (**Fig1c**).

### Optimising serum free expansion of gamma delta T cells

Conventionally, IL-2 is used along with ZOL for expansion, IL-15 is increasingly being explored due to its reported ability to block αβ effector T cells and T-reg cells. Therefore, we tested the impact of IL-15 in the serum-supplemented and serum-free media to investigate its role in the preferential expansion of gamma delta and αβ T cells. Serum-free culturing is one of the goals of cGMP production of therapeutic immune cells. AIM-V (Thermoscientific), OpTimzer(Thermoscientific), SCGM (Lonza), and Immunocult (Stem Cell Technologies) are some of the serum-free media used for testing for their suitability for expanding V gamma 9-V delta 2 T cells (*data not shown*). We observed batch-to-batch consistency with OpTimzer(Thermoscientific) in gamma delta T cells expansion and viability and went ahead with this media for further testing[19]. The fold expansion of the gamma delta T cells was higher in FBS-supplemented media (259+/−55.84-fold, *n=8*) with higher variance (range 42.92 - 472) compared to serum-free media (110+/−29.89-fold, *n=8*) on day 14. However, the samples were more distributed in the FBS cultured cells. The percentage of gamma delta T cells were less variable than αβ T cells (**Fig.02 b,d**). neither serum free conditions nor IL-15 got an impact on the gamma delta T cells to αβ T cell ratio (**Fig.02**).

**Fig 02:**
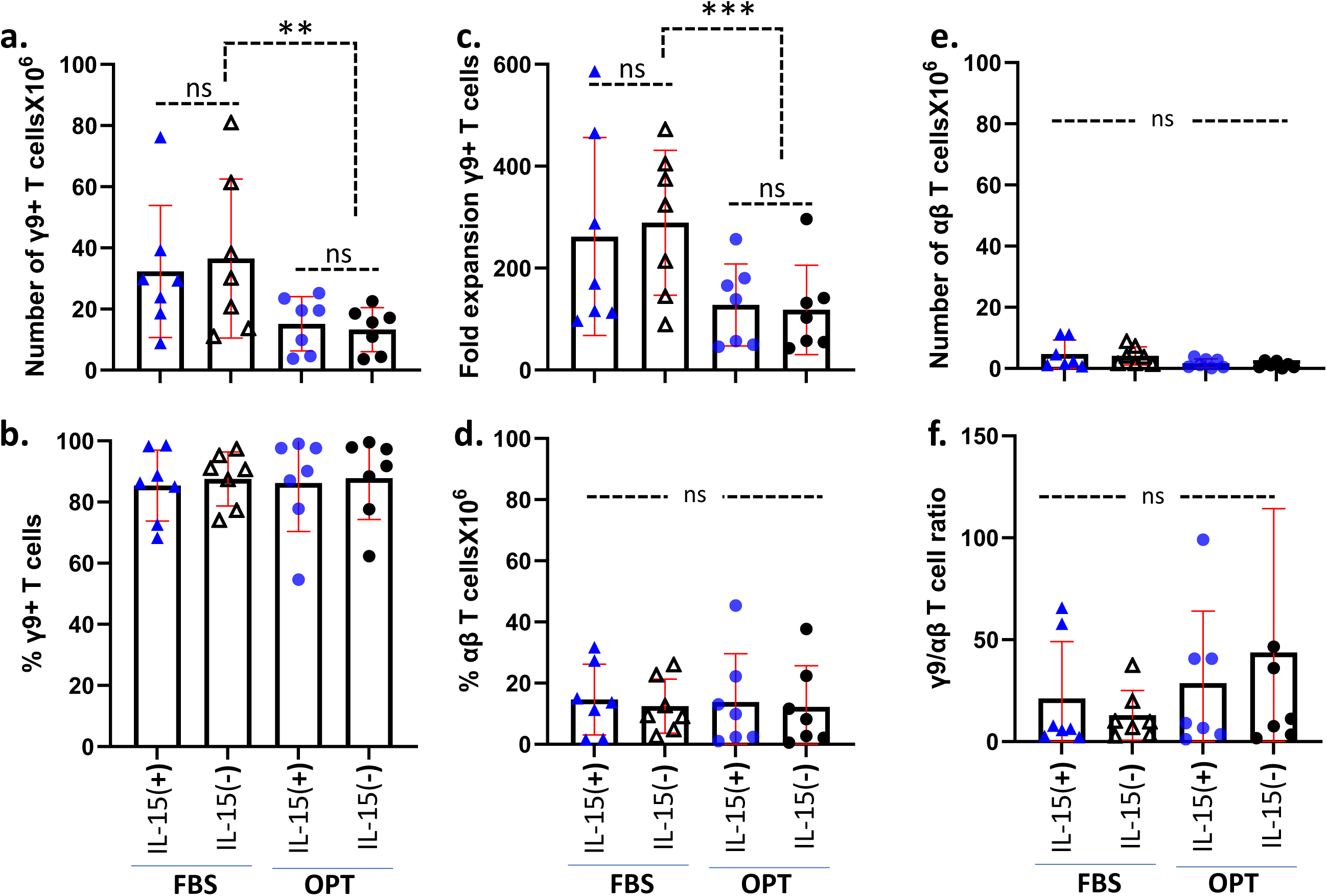
Optimization of gamma delta T cell expansion in serum free media. **a)** Representative flowcytometric graph of T cell populations in the Zol+rhIL-2 expansion culture of PBMC. The phenotype gamma9delta2 T cells on day 14. The percentage of 7-AAD (−) CD3^+^gamma9^+^ T cell populations were analyzed. **b**) The percentage of αβ TCR+ T cells within the live CD3+ populations. **c**) The total number of gamma9delta2 T cells **d**) fold expansion **e**) percentage of gamma delta T cells and **f**) percentage of αβ T cells across the cultures expanded in serum supplemented (RPMI+10%FBS) and serum free (OpTmizer) with or without added IL-15. Each dot represents a normal donor. Data is Mean+/−SEM, n=8, The data is analyzed by student’s t Test. P value is indicated in the figure. The statistical significate is estimated by Student’s T test. * P<0.05, **p<0.00, ***p<0.001.n.s is not significant.

### Kinetic analysis of gamma delta and αβ T cell populations during expansion

Following ZOL activation, there was a logarithmic rise in the percentage of the Vgamma9+ TCR+ population with a concomitant drop in the CD3+αβTCR+ T cells. On day 14, the percentage of gamma delta T cells were enriched to 87.82+/−5.11% (*n=8*) of T cells in Optimizer. The portion of delta2+ within the Vgamma9 compartment is also enriched to more than 90% by day 07, and was increased up to 95% by day 14 (**Fig.3a, Suppl. Fig.02**). The percentage of CD3 neg non-T cell populations (CD19+ B cells, CD14+ Monocytes, CD56 (+) NK cells) were reduced to <3% by day14 (**Fig.3b**). Although gamma delta T cells do not have antigen-specific memory, based on the expression of CD45RA and CD27, they can be divided into naive (CD27 (+) CD45RA (+), central memory (CD27 (+) CD45RA (−), effector memory (CD27 neg CD45RA (−), and terminally differentiated effector memory (CD27(−) CD45RA(+) populations[20]. By day 07, effector memory T cells (CD27^(−)^ CD45RA+) dominated as the central (CD27 (+) CD45RA (−), terminal effector (CD27^(−)^ CD45RA+), and naïve T cell (CD27(+) CD45RA+) population dropped (**Fig.3c**). The fold expansion was to the tune of 75+/−15.41 when 5 samples derived from separate donors expanded in identical conditions were analysed (**Fig.3 d, e, f**). The viability of the gamma delta cells were more than 95% throughout the expansion across 7 donors except for a minor dip in the day 14 mean value (D00: 97.97+/− 3.48; D14: 96.5+/− 1.66) (**Suppl. Fig.03**).

**Figure 03:**
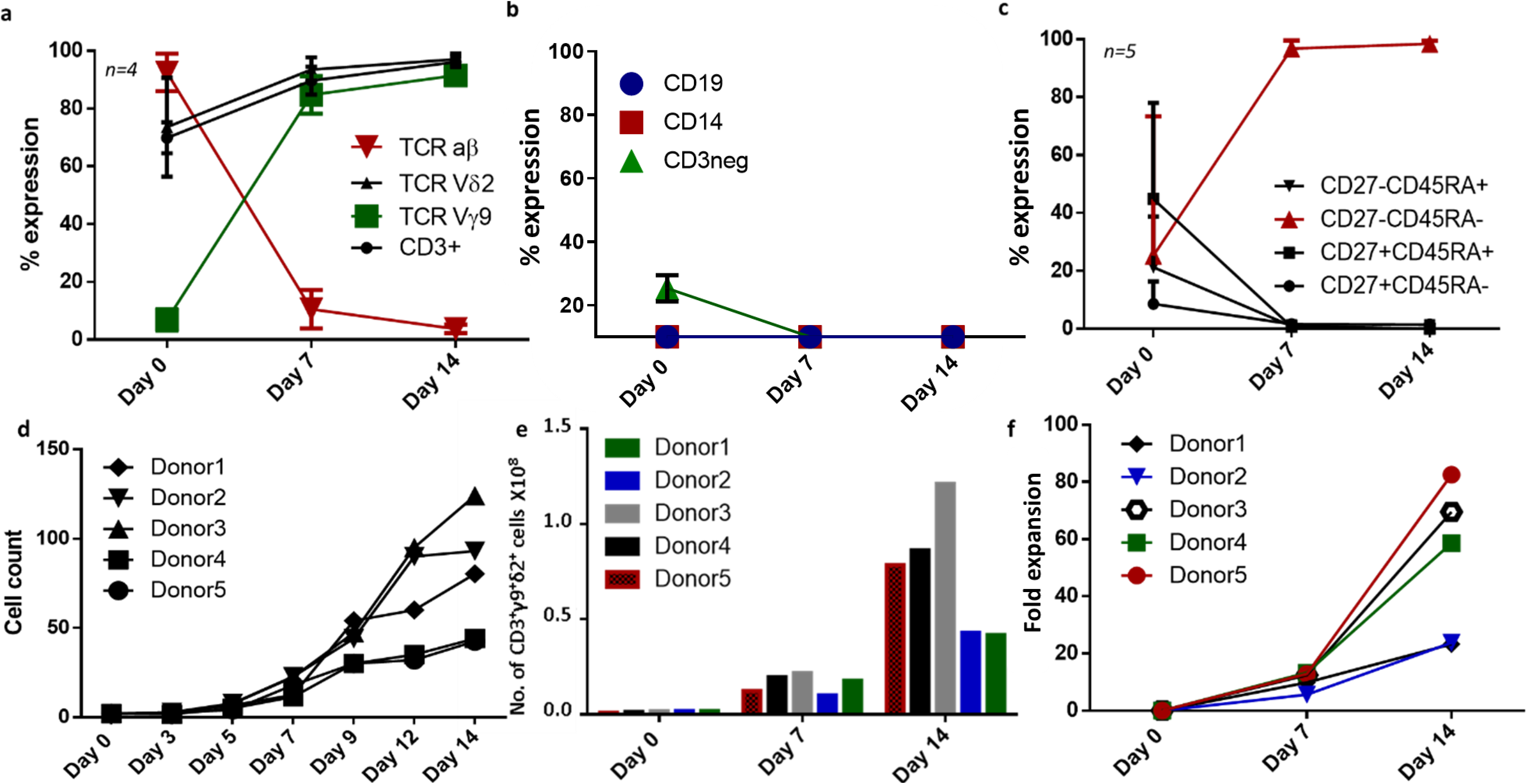
Kinetics of gamma delta T cell expansion in the ZA+IL2 activated culture of PBMC. **a**. Percentage expression of the indicated populations as measured by flow cytometry at day0, day7, and day14. mean+/−SEM n=4. **b**. Percentage expression of non-lineage populations such as total live CD3neg population, CD14+ monocytes, and CD19+ B cells. mean+/−SEM from n=7. **c.** Percentage expression of naïve (CD45RA+CD27+), effector memory (CD45RA-CD27-), central memory (CD45RA-CD2+), terminally differentiated (CD45RA+CD27-). The data are mean+/−sem, n=5. **d.** A total number of cells in the culture of an individual donor over the indicated dates. n=5 **e**. The number of CD3+gamma9+delta2+ cells calculated from the total number of cells in the culture, n=5 **f.** Fold expansion of CD3+gamma9+delta2+ cells calculated against baseline numbers of the same. n=5. The statistical significate is estimated by Student’s T test. * P<0.05, **p<0.00, ***p<0.001.n.s is not significant.

NKG2D+ population within the CD3+Vgamma9+ compartment was enriched during expansion (D00: 75.28+/−7.48, D14: 93.76+/−1.55; n=5, **Suppl.Fig.04a**). NKG2D expression on day 07 and day 14 was significantly high on Vgamma9+delta2+ population indicative of its potential antitumor functions (**Fig.04 a, b**). CD69 is an early activation marker often associated with tissue residency in the T cells. By day 14, over 90% of gamma9+ cells were CD69+NKG2D+ T cells, representing activated cells with strong cytotoxic potency (**Suppl.Fig.4b**).

**Fig. 04:**
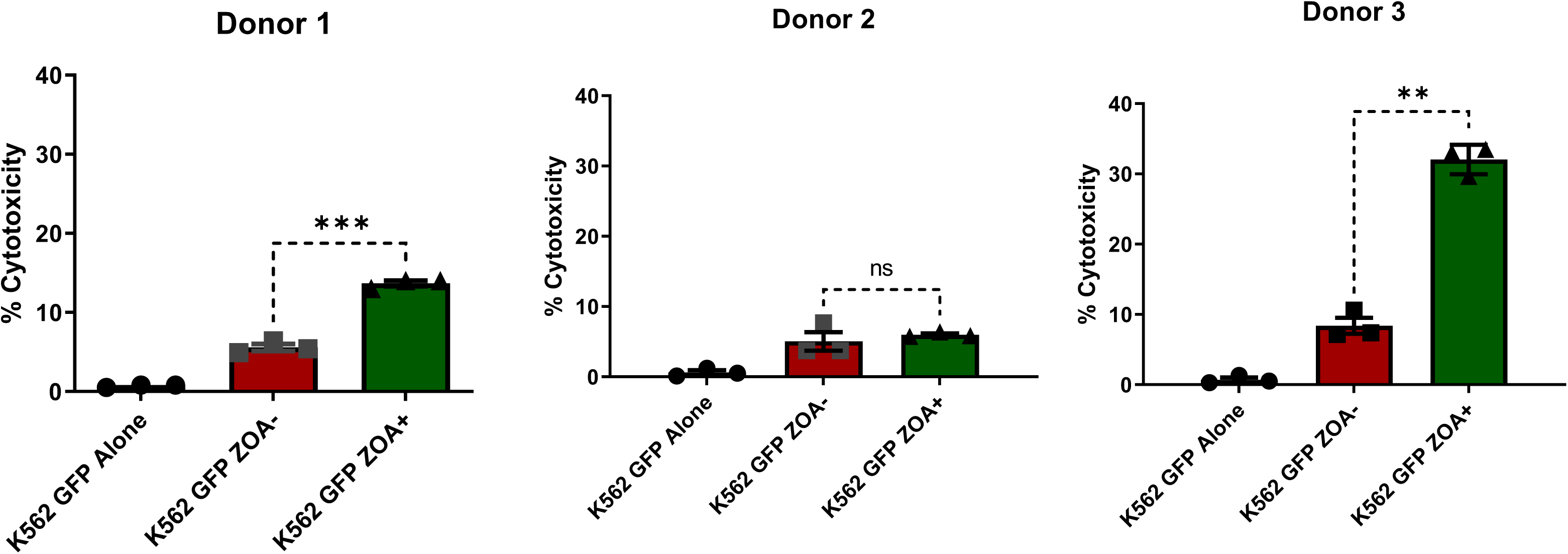
Zoledronic acid pretreatment of tumor cells enhance the cytotoxicity of Vgamma9+Vdelta2+ T cells. The Vgamma9+Vdelta2+ T cells were harvested on day 14 (materials and method) and was co-cultured with K562-GFP cells (1:1). K562-GFP cells were pretreated for 6 hrs with Zoledronic Acid (5µM) before washing and plating for the co-culture. The Vgamma9+Vdelta2+ T cells were harvested after 4 hrs of co-culture and the percentage cytotoxicity was calculated as per the suppl.fig.5a. fig a, b and c. The data is represented as mean +/−SEM (r=3). The statistical significate is estimated by Student’s T test. * P<0.05, **p<0.00, ***p<0.001

### Pre-treatment of target tumor cells with Zoledronic Acid impacts the tumor-toxicity V gamma 9-V delta 2 T cells in a donor-dependent manner

A crucial parameter that gauges the therapeutic potential of the effector immune cells is its ability to lyse tumor cells with reduced off-tumor toxicity. We have confirmed that these gamma delta T cells lyse established leukemic cell lines K562-GFP (CML cell line) in various effector-to-target ratios as revealed by a flow-based assay developed in our lab. The percentage cytotoxicity was 15.35 +/− 2.71 at 1:1 ratio and 41.60 +/− 2.415 (Mean+/−SEM, n=2) (**Suppl. Fig.05**). Pre-treatment of tumor cells with ZA was reported to enhance the tumor toxicity of V gamma 9-V delta 2 T cells[21–23]. Therefore, we have pre-treated the tumor cells (K562) with 5µM of ZOL for 6hrs followed by washing and co-culturing with gamma delta T cells at a 1:1 ratio for 4 hours to measure the percentage of immunogenic cytotoxicity. We observed that in gamma delta T cells derived from 2 out of 3 donors, ZOL pretreatment significantly enhanced the cytolysis of the K562 cells (donor 1 untreated 5.57+/−0.44% to treated 13.66+/−0.33%; p=0.009 and donor 3 untreated 8.383+/−1.12% to treated 32.06+/−1.21%; p=0.003). However, for the second donor, there was no change (p=0.580), perhaps indicating donor-to-donor variations in the configuration of the tumor sensing receptor in gamma delta T cells (untreated 5.04+/−1.30 treated 5.99+/−0.19) (**Fig.04**). The mean difference and fold enhancement in cytotoxicity due to ZOL pretreatment is 10.88+/−6.68 and 2.48+/−0.76 respectively (**Suppl.Fig.06**).

The IFNgamma production and CD107 degranulation strongly indicate activated gamma delta T cells poised for cytotoxicity. Therefore, the expanded gamma delta T cells were co-cultured with ZOL(5μM) pre-treated K562 cells and the IFNgamma production and CD107a degranulation was measured by flow cytometry after 6 Hrs. Interestingly, ZOL pre-treatment could enhance the IFNgamma secretion only to a less than significant scale across these experiments. CD107 degranulation was found to be enhanced significantly upon tumor cell encounter (**Fig. 05a, b**). We observed that gamma delta T cells expanded in serum free media produced more IL-10 with no significant change in the production of TGF β. Furthermore, a significant enhancement in the CD16 expression the gamma delta T cells expanded in the serum free expanded cells was observed. However, expression of cytotoxicity markers such as Fas L, NKp30, NKp44 and TRIAL appears to remain similar across serum free media and optimizer (**Suppl.Fig 07 and 08**).

**Fig. 05:**
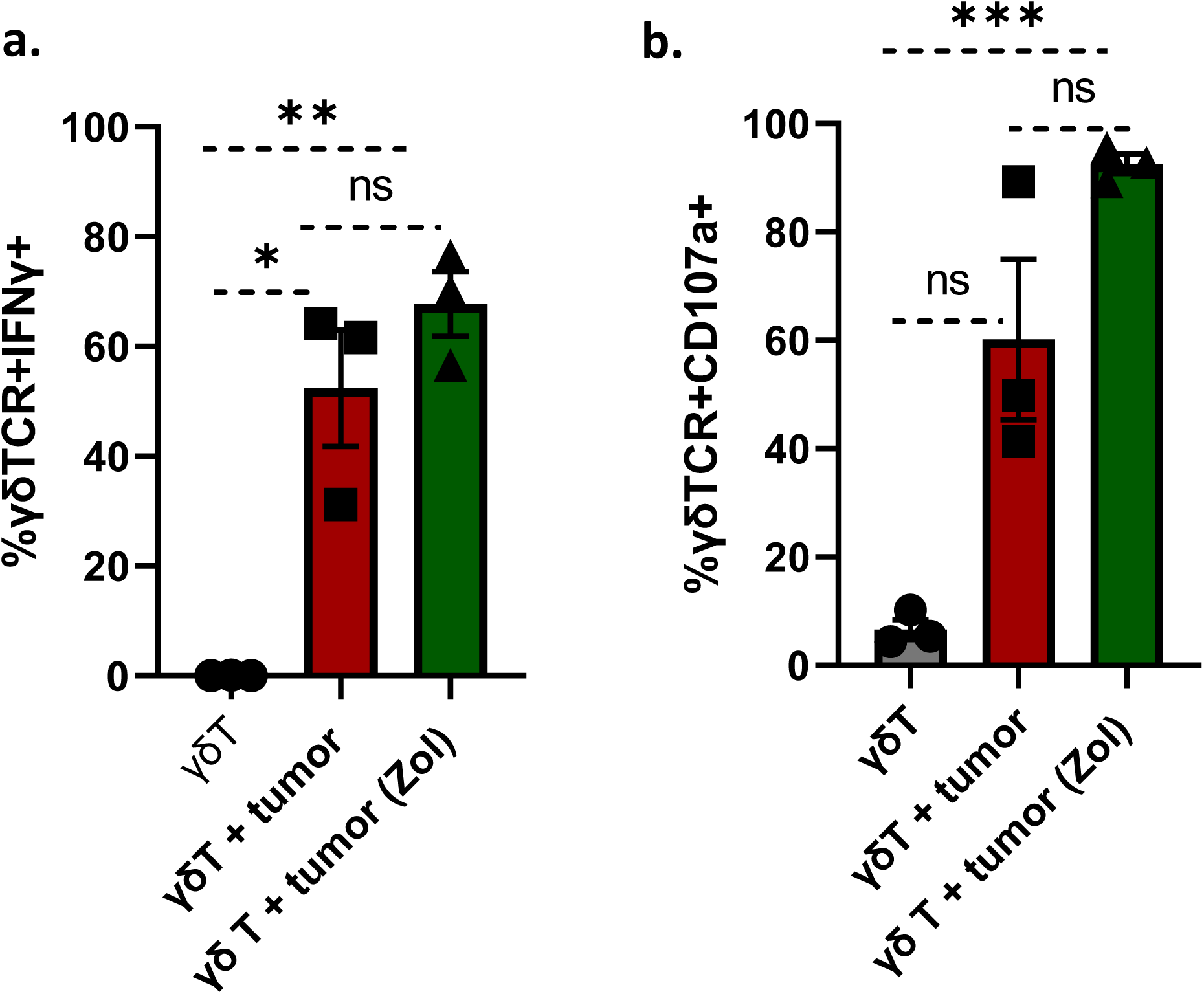
IFNgamma and CD107a from gamma delta T cells: a,b. IFNgamma and CD107a is assed from live+gamma delta TCR+ populations. The data is represented as mean +/−SEM (n=3, r=3). The statistical significate is estimated by Student’s T test. * P<0.05, **p<0.00, ***p<0.001

### Engineering of V gamma 9-V delta 2 T cells expanded in serum-free media

Genetic modification to enhance the therapeutic index is a primary downstream application of the therapeutic expansion of immune cells. Towards that end, we have optimized the lentiviral transduction of V gamma 9-V delta 2 T cells. Although promoters play a crucial role in driving transgene expression in cell and gene therapy, **t**he outcome of promoters varies across the host cell. Therefore, we have compared the transduction efficiency of two commonly used promoters −CMV and EF1α-often used in driving therapeutic transgenes. The V gamma 9-V delta 2 T cells transduced lentivirally by pEF1α-GFP was a gift from Dr. Trent Spencer, Emory University and pCMV-GFP was purchased from addgene (*addgene plasmid#36083*) at 80 MOIs in serum-free media (OpTmizer, Thermo scientific) with gamma delta T cells plated at a density of 0.5e6/ml. Calcium Phosphate, a cGMP-compatible reagent, was used to assist transfection. The transduction efficiency is monitored by GFP expression every 2 days until day 14. The transduction efficiencies in pEF1α (68.10+/−9.09) and pCMV-GFP (39.09+/−8.94% n=3, n.s.) indicates the utility of both promoters for driving transgene expression stably in gamma delta T cells. However, the viability of the gamma delta T cells expressing CMV promoter was much higher suggesting possible advantage of using CMV promoter for CAR transduction in gamma delta T cells (pCMV 84.01% +/− 6.465 n=3; pEF1α 19.17% +/− 5.190 n=3) (**Fig.06**). Together, our data suggests the feasibility of using CMV promoter for engineering gamma delta T cells for adoptive immunotherapy applications.

**Fig. 06:**
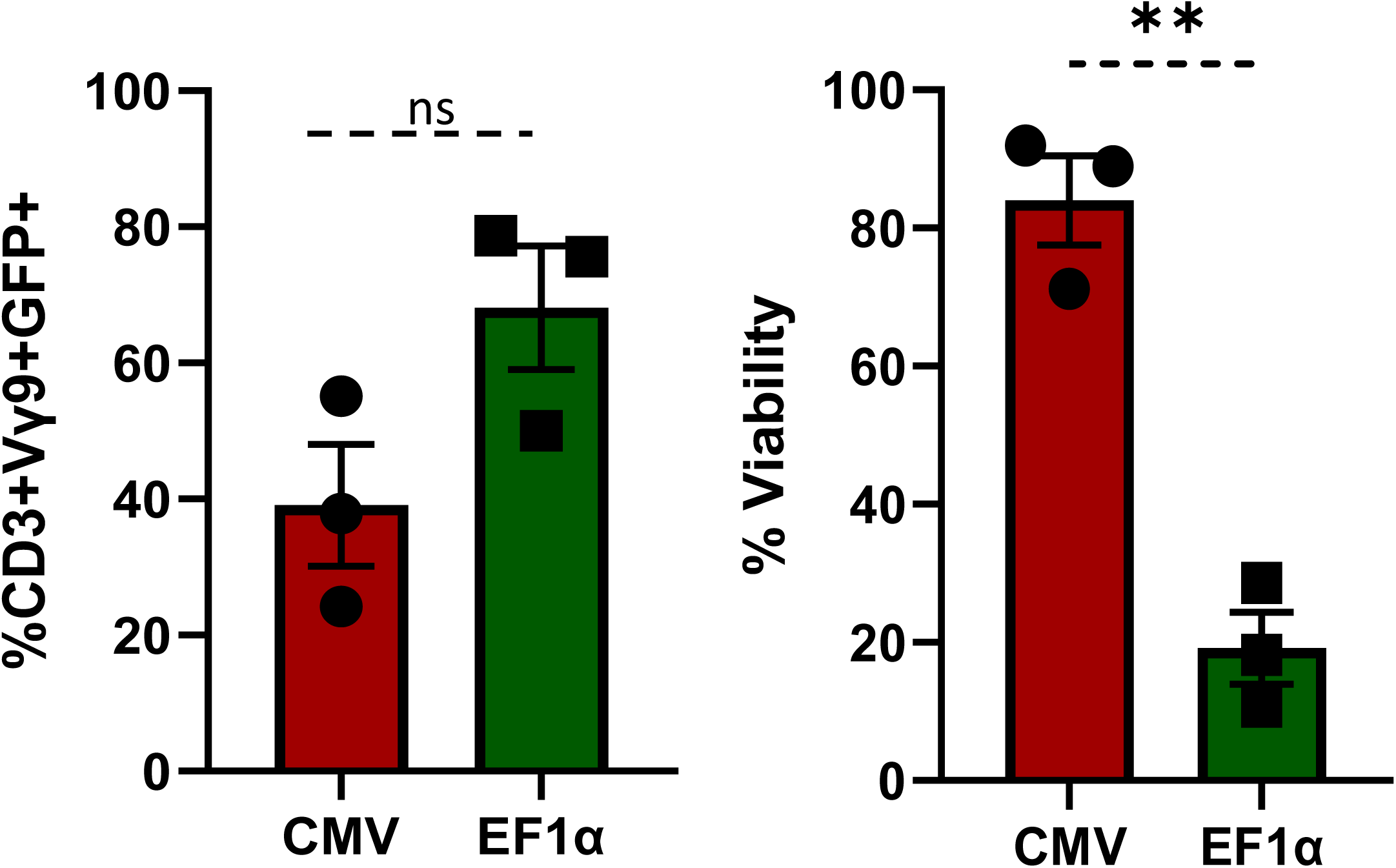
Lentiviral transduction of gamma delta T cells: a. The strength of CMV and EF1α was quantified against the flow cytometric expression of the driven GFP transgene in Vdelta2+ subsets. a) The percentage viability as measured by 7AAD+ cells in the Vgamma9+Vdelta2+ T cells 28 hrs after lentiviral transduction. B) The transduction efficiency of GFP vector on Vdelta2+ subsets in the above experiments. The percentage expression of CD3+7AAD (neg) Vgamma9+Vdelta2+ T cells. The data is represented as mean +/−SEM (n=3, r=3). The statistical significate is estimated by Student’s T test. * P<0.05, **p<0.00, ***p<0.001

## Discussion

Activation of PBMCs with ZOL (an FDA-approved drug for osteoporosis) and IL-2 is a standard method for expanding human V gamma 9-V delta 2 T cells^24,25^. However, a streamlined protocol for serum-free expansion must be optimized. Apart from purity, the cytokine requirement for expansion in serum-free / xeno-free media, reduced cytotoxicity, prolonged expansion, reduced transduction with lentiviral transduction, etc., are not adequately addressed.

Our results indicated peripheral blood as a better starting source for delta2 (+) T cells, while cord blood had a higher percentage of delta2 (−) population in the gamma9(+) compartment. Although delta1 (+) and delta3 (+) T cells are the other considerable populations within the gamma9(+) T cells, the identity of other populations needs to be investigated further[24].Serum-free expansion reduced the overall yield of gamma delta T cells than serum-sufficient media. However, purity and inter-donor congruence in fold expansion is considerably enhanced. In contrast to the previous protocols, we achieved higher gamma delta T cell purity as early as day 07 with the enrichment of delta2 (+) CD45RA^neg^CD27^neg^ effector cells with antitumor functions on day 14. The αβ T cells and non-lineage cells were less than 2% on day 14, permitting the depletion with reduced amount of antibodies in the allogenic settings or skipping the depletion step altogether in the autologous settings. The role of IL-15 in the expansion of gamma delta T cells appears variable across the donors indicating, a need to further investigate the molecular basis of IL-15R signalling[25]. This distinguishes our studies from the previous reports of enhancement in proliferation and cytotoxicity of gamma delta T cells upon expansion in the presence of IL-15^16^. Nonetheless, the higher purity of gamma delta T cells in our protocol can be a trades off against the higher yield reported in the previous protocols[26,27]. The composition of IFNgamma+ gamma delta T cells with antitumor functions and IL-17+ gamma delta T cells with protumor roles is decisive while characterizing the therapeutic gamma delta T cells. Furthermore, the mutually contradictory functions of IL-2 and IL-15 in the expansion of protumor IL-17+ gamma delta T cells deserves further characterization of IFNgamma and IL-17 populations in our product[28,29].

Allogenic αβ TCR-positive cells must be depleted before transfusion to prevent outgrowth and potential GvHD. Because of the higher purity on day 14, αβ T cell depletion can be achieved by relatively reduced amount of antibody cocktail and potentially save the cells lost during the depletion, cutting the cost of the therapy. Currently, from 1 million peripheral blood mononuclear cells, we can expand up to 75+/−15.41 million gamma delta T cells in the serum-free media. However, this can indicate compromised the cell number against the higher purity possibly due to lack of growth factors from the αβ T cells[30,31]. Further enrichment of the serum-free media with additional growth factors/cytokines can potentially enhance the proliferation and function of gamma delta T cells[32]. In fact, with alteration in the media changing pattern, we could further enhance the purity of gamma delta T cells than that observed in Fig.02 *albeit* with a reduction in the yield (Fig.03 onwards).

Since the purity of gamma delta T cells at day 07 is approaching industry standard, we interrogated the expression of NKG2D - which predicts antitumor functions - on Day 07 and Day 14 and observed it to be comparable. Since longer *in vitro* expansion can potentially cause exhaustion, our results demand further experiments to compare the cytotoxic capacity on day 07 and day 14. Although gamma delta T cell therapy was well tolerated in clinical trials, the antitumor functions were suboptimal to standard of care, necessitating further modification. Importantly, ZOL pretreatment was found to enhance the tumor toxicity of gamma delta T cells in a donor-dependent manner[33]. Recently, ATRA and Ibrutinib pre-treatment were found to augment the antitumor functions of V gamma 9-V delta 2 T cells[34]. This opens possibilities to potentially combine such small molecules in sublethal concentrations to enhance the tumor lysis capacity of V gamma 9-V delta 2 T cells.

Lymphocyte engineering with CARs is another approach to improvise the antitumor functions of gamma delta T cells. However, gene delivery to gamma delta T cells is more challenging than the conventional αβ T cells due to increased susceptibility to cell death and reduced expression of low-density lipoprotein receptor (LDL-R) receptors. Therefore, several transductions at a higher multiplicity of infection (MOI) are often required[35]. Our protocol is optimized for single transduction - at 80 MOI, using Calcium Phosphate as transfection reagent - an advancement compared to established protocols.

Affordability of the cell therapeutics is the crucial factor that is limiting the wider adoption across the health systems even in developed nations. Automated batch production of an off-the-shelf cell therapy within in shorter period can reduce the price by an impressive margin (30770285, 33674239). Our protocol can cut the cost owing to the production of an unmodified off-the-shelf product with improved purity that can be achieved even in a shorter span of time. The reduced logistical complexity of gamma delta T cells may further reduce the cost. Although the reduced yield of gamma delta T cells can be the limitation of this approach, possibility to rationally combine with small molecules and antibodies can be experimented to incrementally achieve the window of therapeutic response without hiking the cost.

Our protocol uses cGMP-compatible components such as FDA-approved drugs - ZOL, and IL-2 in serum-free media to produce V gamma 9-V delta 2 T cells with higher purity and diminished inter-donor variability. Our results also suggest for reducing the time of expansion because of the favourable enrichment of delta2(+) NKG2D+ T cells. Furthermore, this protocol eliminates the requirement for αβ T cell depletion for off-the-shelf therapy. V gamma 9-V delta 2 T cells thus expanded lysed the tumor cells *in vitro* and can be further enhanced by pre-treatment of tumor cells with ZOL. Higher purity, reduced expansion time, and the possibility of combining ZOL can compensate for the reduced cell number and enhance the overall efficacy. Remarkably, our protocol allows engineering V gamma 9-V delta 2 T cells with single transduction. It is tempting to speculate that this protocol may be further fine-tuned for the automated and rapid production of V gamma 9-V delta 2 CAR T cells to potentially reduce the cost and enhance the antitumor functions of CAR T cells.

## Acknowledgments

We acknowledge the funding support from start-up fund from Center for Stem Cell Research and Ramalingaswamy Re-entry Fellowship (No. BT/RLF/Re-entry/35/2016) to Dr. Sunil Martin. We thank Dr. Shaji Ramachandran Velayudhan for core facility support and Dr. Sonum Pandey for data archiving.

## Supplementary Figure Legends

**Suppl.Fig.01: Workflow for expansion of Vgamma9+Vdelta2+ T cells**. Peripheral blood derived mononuclear cells (1 million/mL) is plated in 24 well plate and pre-treated with 5μM of Zoledronic Acid and recombinant human IL-2 (1000 IU) with or without IL-15 (for certain experiments) in CTS OpTmizer as per the methodology. The cells were washed and supplemented with media and cytokines on day 03, 05,07,09,11 and 13. Cells were harvested on day 00, 07 and 14 for counting, phenotyping and functional analysis.

**Suppl.Fig.02: The flowcytometric analysis of Vgamma9+Vdelta2+ T cells during expansion**. The Vgamma9+ and Vgamma9+Vdelta2+ populations are monitored within the CD3+7AAD (neg) live cells on 0th, 7th and 14^th^ day. The data is a representative figure from Fig.03.

**Suppl.Fig.03: Viability of the *in vitro* expanded Vgamma9+Vdelta2+ T cells.** The percentage viability of the Vgamma9+Vdelta2+ T cells in the gamma delta T cell expansion experiments as measured by 7-AAD staining within the lymphocyte gate. The data is represented as mean+/−SEM across 07 donors on day 00 and day14.

**Suppl.Fig.04: NKG2D+ expression during the expansion of gamma delta T cells. a.** The expression of NKG2D alone within the Live+CD3+ Vgamma9+ cells on day 00 and day 14 as measured by flowcytometry. The data is represented as mean +/−SEM (n=5). **b**. The percentage expression of NKG2D and CD69 within the Live+CD3+ Vgamma9+ and the Live+CD3+ Vgamma9+ Vdelta2+ gate. The data is represented as mean +/−SEM (n=4). The statistical significate is estimated by Student’s T test. * P<0.05, **p<0.00, ***p<0.001.

**Suppl.Fig.05: Cytotoxicity assay set up for gamma delta T cells a)** The protocol for the flow-based cytotoxicity assay. The difference between the calculated 7-AAD positivity from 4th hr and 0th hr is indicated as percentage cytotoxicity **b)** The percentage cytotoxicity of gamma delta T cells at two effectors to target ratios (1:1 and 1:5). The data is represented as mean +/−SEM (n=2). The statistical significate is estimated by Student’s T test. * P<0.05, **p<0.00, ***p<0.001.n.s is not significant.

**Suppl.Fig.06. ZOL enhances the sensitivity of the tumor cells to gamma9delta2 T cells differentially.** The delta (a) and fold expansion (b) data calculated from Fig.04. Each dot represents the delta and fold expansion for individual donors.

**Suppl.Fig.07. Production of anti-inflammatory cytokine from gamma delta T cells:** a. TGF-β and b. IL-10 were measured by ELISA from the media of gamma delta T co-cultured with tumor cells. (n=3, Mean+/−SEM). * p ≤ 0.05.

**Suppl.Fig.08. The expression of cytotoxic markers (CD16. FasL, NKp30, NKp44 and TRIAL).** The expression of cytotoxic markers (CD16. FasL, NKp30, NKp44 and TRIAL) were measured by flowcytometry on day 0, day 7 and day 14 of the culture. **a**. Representative FACS plots (day 14). b. CD16 **c**. Fas ligand d, NKp30 **e**. NKp44 and **f.** TRIAL were measured from gamma delta T cells expanded in serum-free and serum supplemented media (n=3, Mean+/−SEM). ** p ≤ 0.01.

