## Supplementary figures and images for "Serum free expansion and transduction of human V gamma 9-V delta 2 T cells for adoptive immunotherapy"

### Supplementary Figures

Suppl.Fig.01

**Suppl.Fig.1**

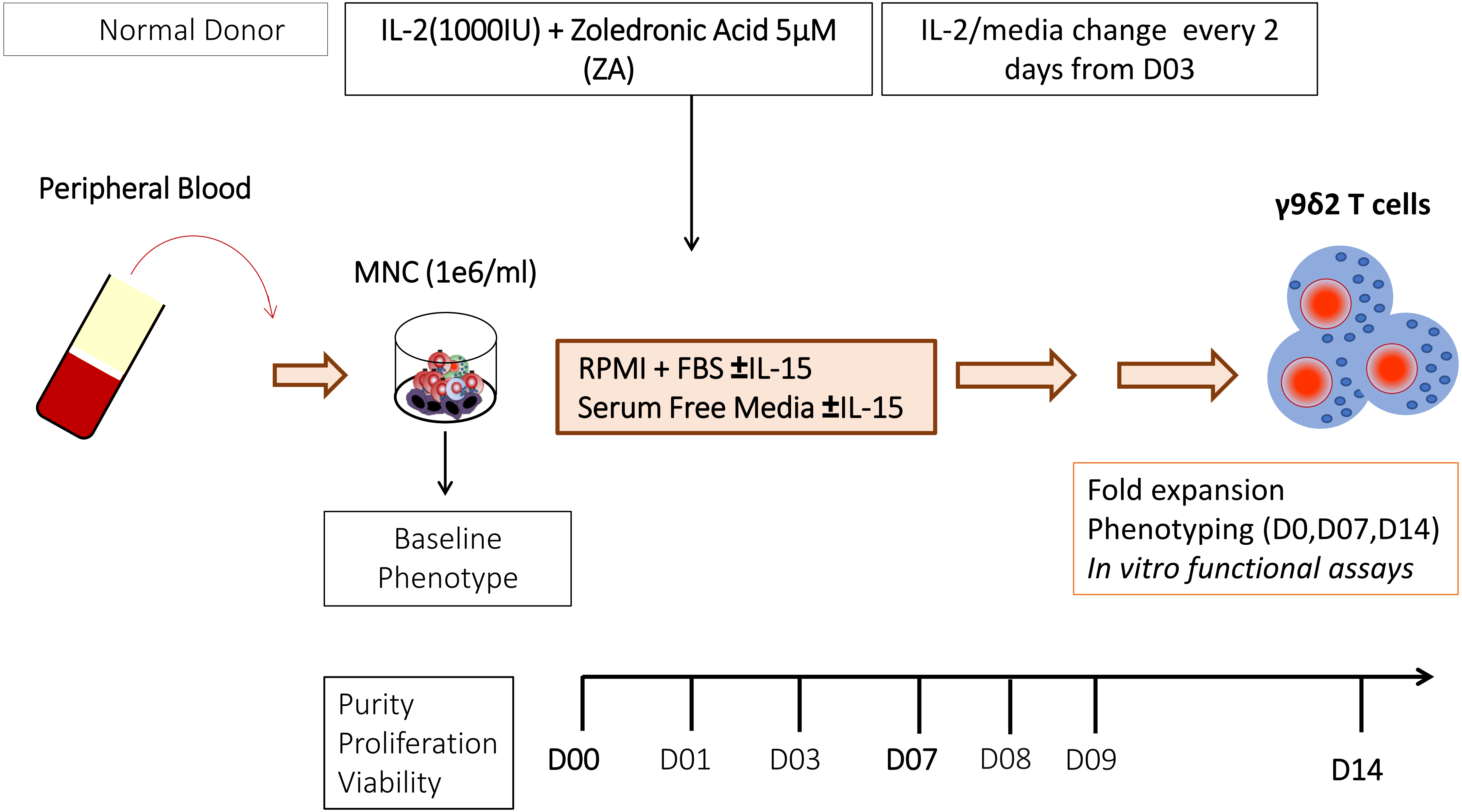

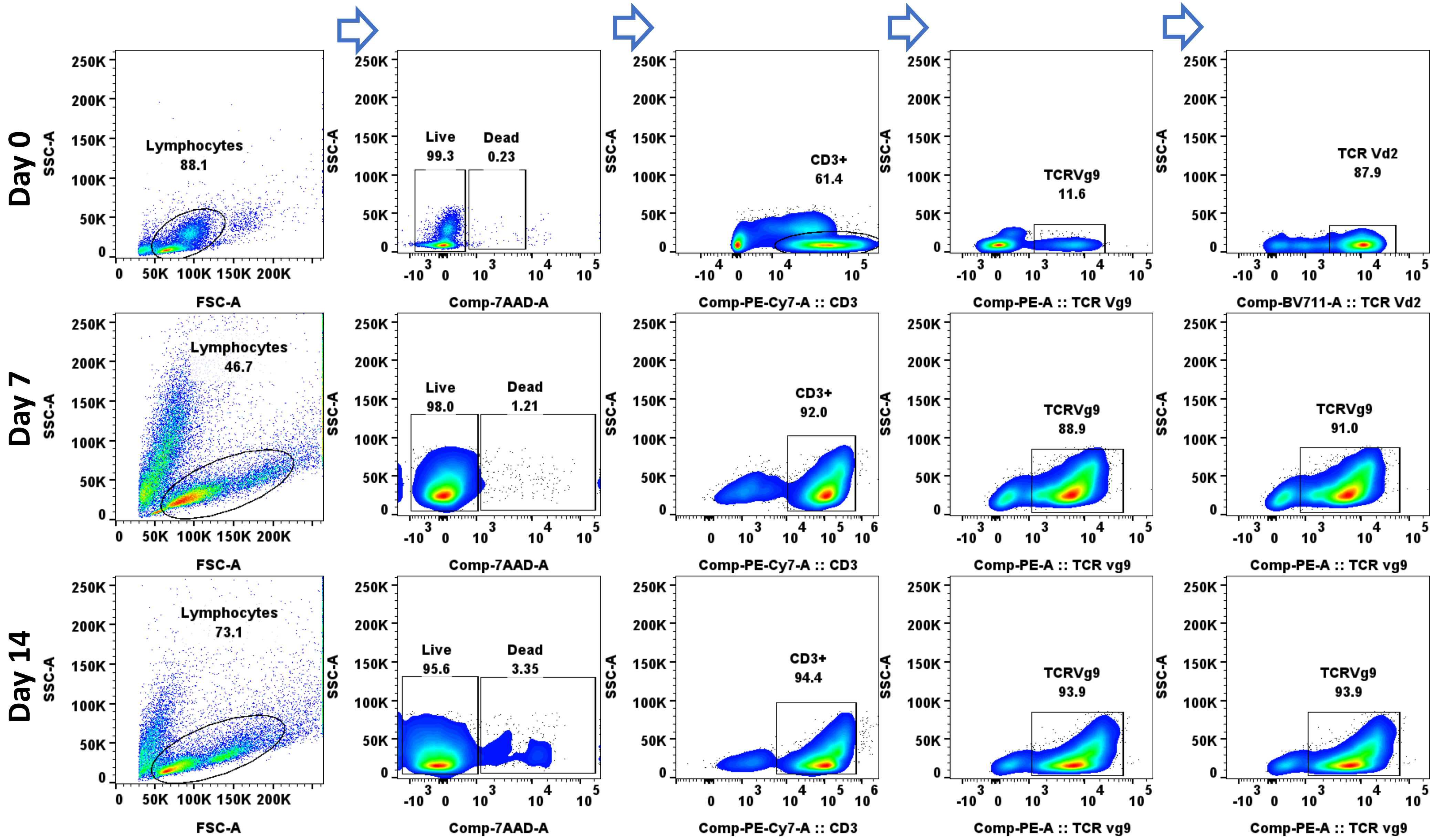

Suppl.Fig.03

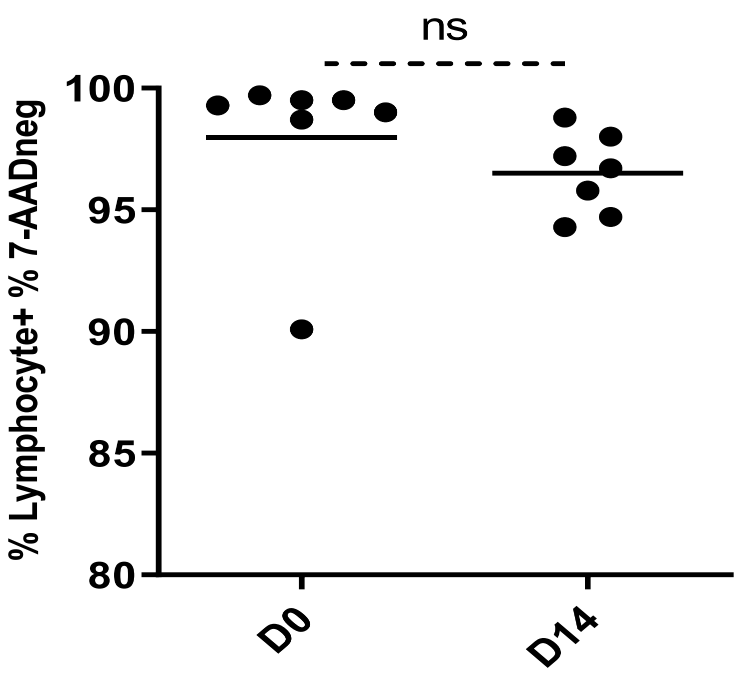

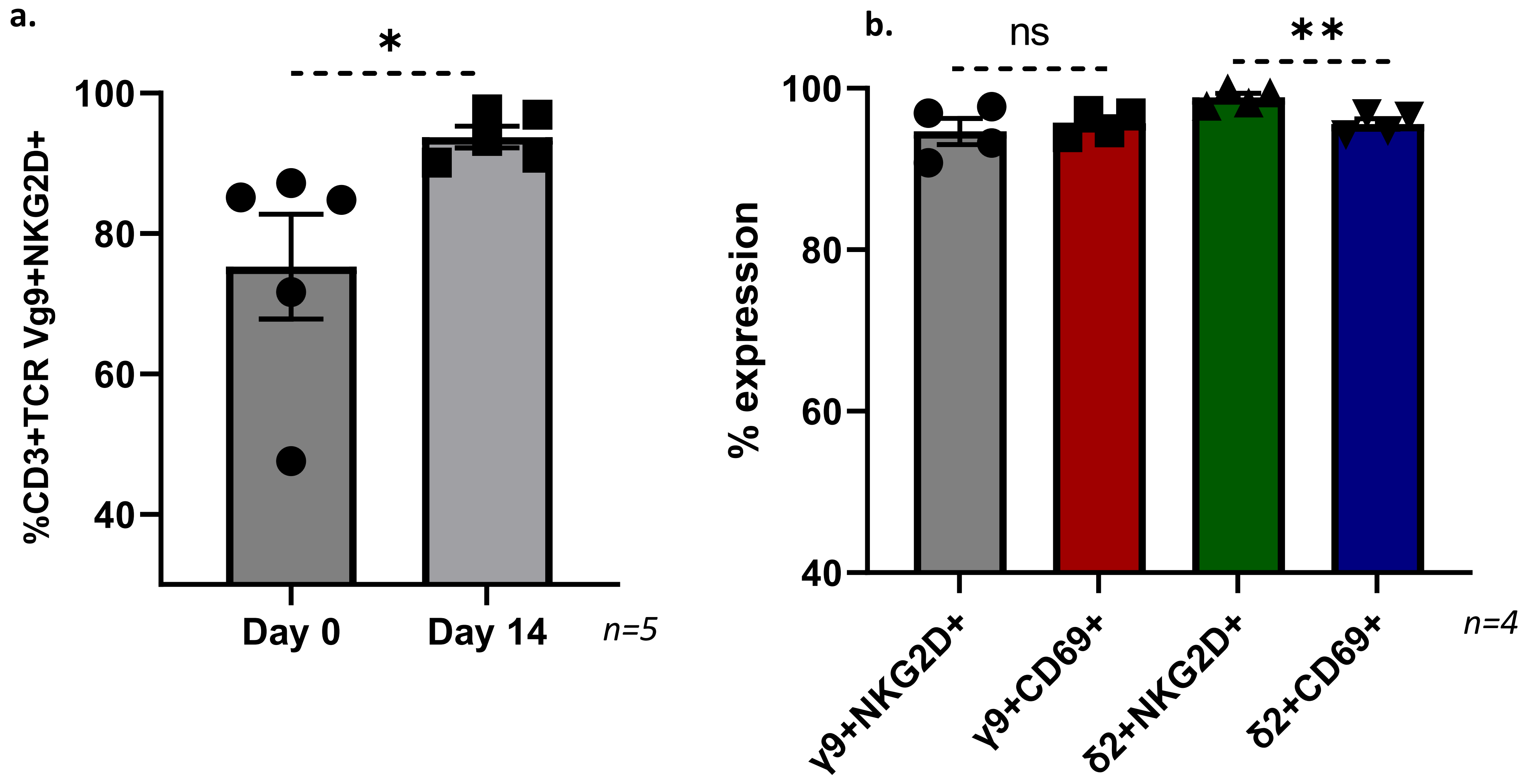

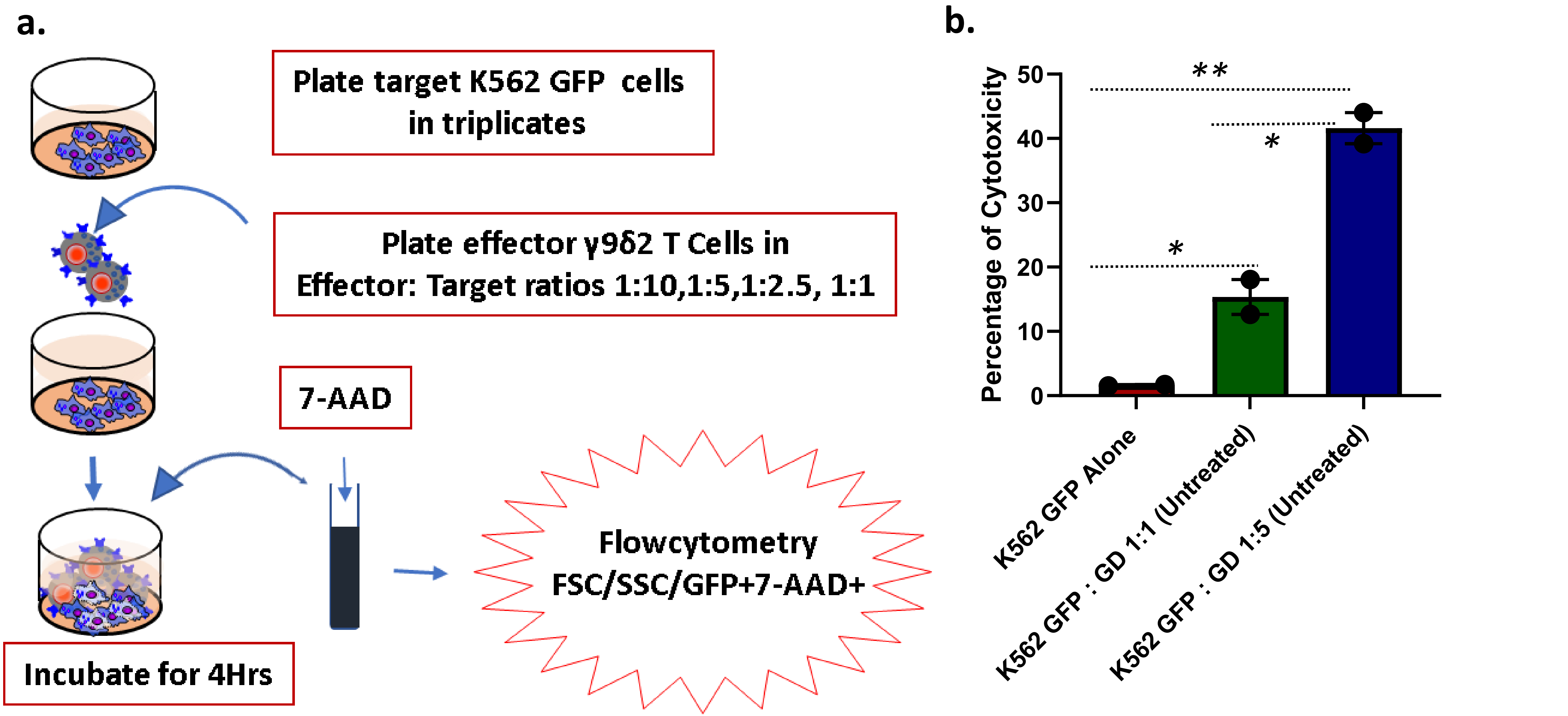

Suppl.Fig.06

a.

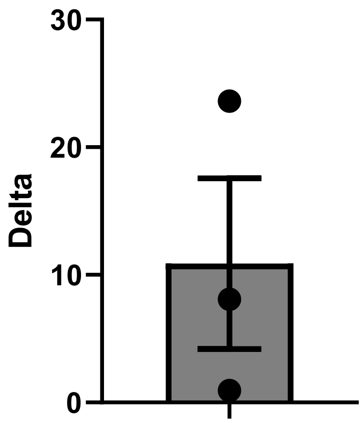


Mean±SEM n=3
10.88±6.68

b.

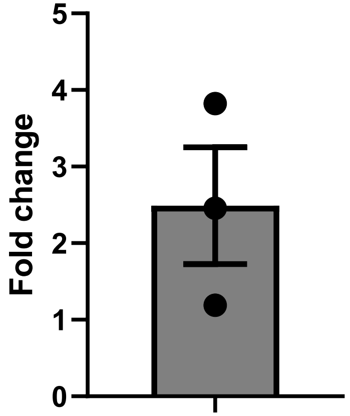


Mean±SEM n=3
2.48±0.76

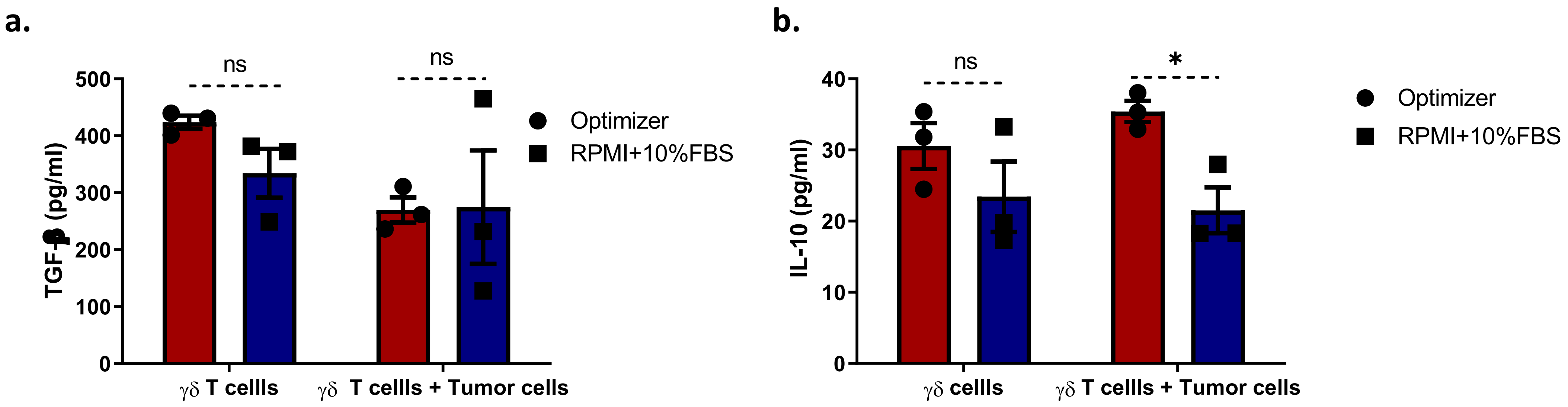

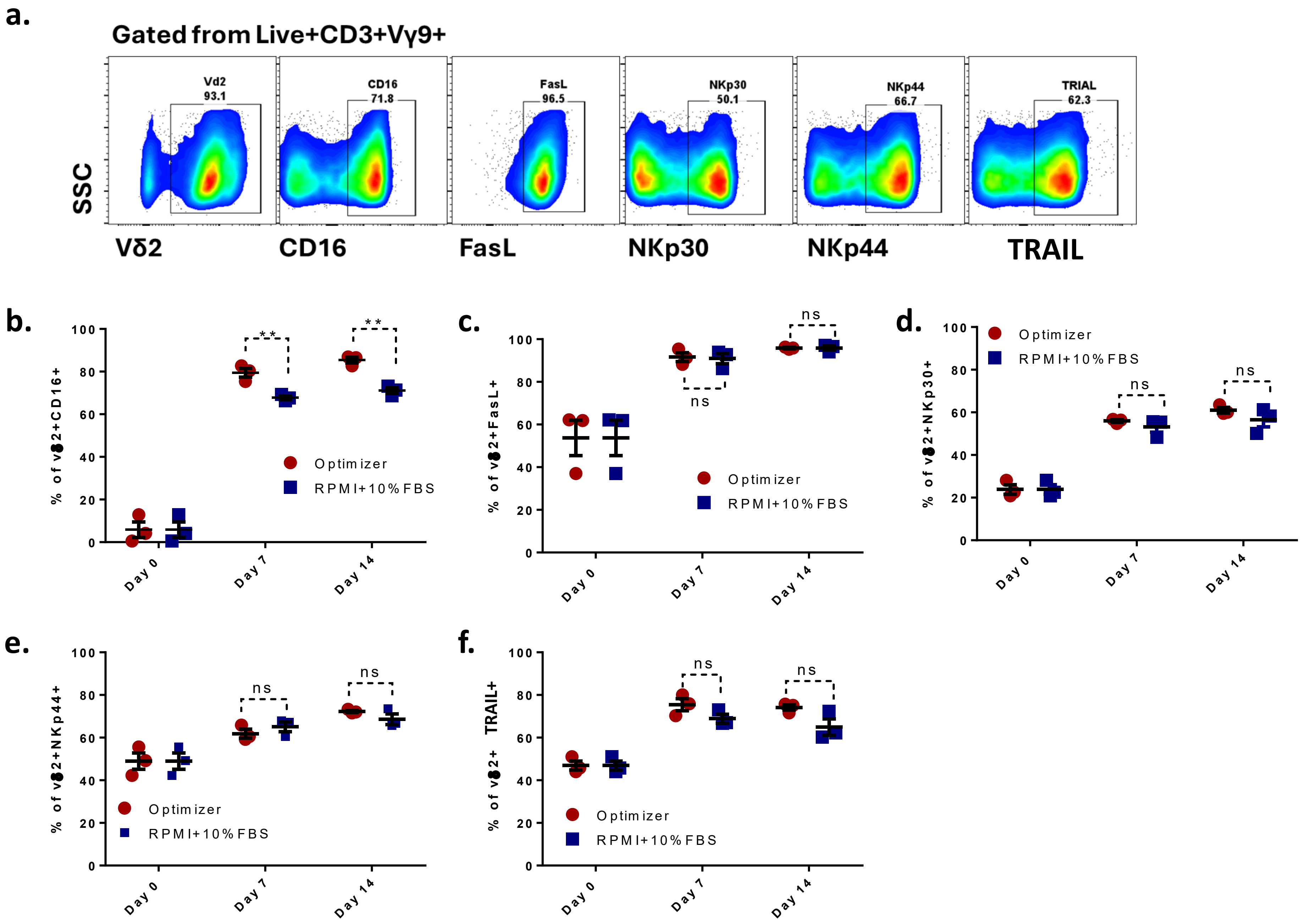
